# Defensive shimmering responses in *Apis dorsata* are triggered by dark stimuli moving against a bright background

**DOI:** 10.1101/2022.02.21.481276

**Authors:** Sajesh Vijayan, Eric J Warrant, Hema Somanathan

## Abstract

Giant honeybees, including the open-nesting Asian giant honeybee *Apis dorsata*, display a spectacular collective defence behaviour – known as “shimmering” – against predators, which is characterised by travelling waves generated by individual bees flipping their abdomens in a coordinated and sequential manner across the bee curtain. We examined if shimmering is visually-mediated by presenting moving stimuli of varying sizes and contrasts to the background (dark or light) in bright and dim ambient light conditions. Shimmering was strongest under bright ambient light, and its strength declined under dim-light in this facultatively nocturnal bee. *A. dorsata* shimmered only when presented with the darkest stimulus against a light background, but not when this condition was reversed (light stimulus against dark background). This response did not attenuate with repeated exposure to the stimuli, suggesting that shimmering behaviour does not undergo habituation. We suggest that this is an effective anti-predatory strategy in open-nesting *A. dorsata* colonies which are exposed to high ambient light, as flying predators are more easily detected when they appear as dark moving objects against a bright sky. Moreover, the stimulus detection threshold (smallest visual angular size) is much smaller in this anti-predatory context (1.6° - 3.4°) than in the context of foraging (5.7°), indicating that ecological context affects visual detection threshold.

## Introduction

Honeybee colonies consist of several thousand individuals and brood cells, as well as abundant stores of honey and pollen that must be guarded from predators (Fuchs and Tautz, 2011; Seeley, 1995). The giant honeybee *Apis dorsata* is a tropical open-nesting species with colonies occupying cliff faces, tall trees or man-made structures such as water tanks and window ledges (Misra et al., 2016; Neupane et al., 2013; Roy et al., 2011; Tan et al., 1997; Thomas et al., 2009). Honeybee species exhibit a wide array of anti-predatory behaviours (Fuchs and Tautz, 2011; Phiancharoen et al., 2011; Seeley et al., 1982), and shimmering behaviour reported in the giant honeybees is the first line of defence against aerial flying predators such as birds and hornets which are their major predators (*A. dorsata*: (Kastberger et al., 2011), *A. laboriosa*: (Batra, 1996)). This behaviour is visualised as a dark travelling wave on the comb surface, and can precede a more aggressive collective stinging response (Seeley et al., 1982; Kastberger and Sharma, 2000; Kastberger et al., 2011; Wongsiri et al., 1997).

Shimmering in *A. dorsata* arises from ‘flickering’ behaviour, a form of dorsoventral abdomen-flipping that occurs stochastically (Weihmann et al., 2012). Once initiated at trigger centres on the bee curtain (Kastberger et al., 2010a; Schmelzer and Kastberger, 2009), the shimmering wave follows the path of the flying intruder (Kastberger et al., 2010a). More trigger centres can subsequently form if the intruder persists, as quiescent bees from other regions on the comb initiate shimmering, leading to progressively stronger and larger displays (Weihmann et al., 2012). The waves also accelerate at a few trigger centres that detect approaching waves from further away (Kastberger et al., 2012; Kastberger et al., 2014). Shimmering confuses potential predators, and serves as a multi-modal alerting signal for nestmates using mechanoreceptive cues (Kastberger et al., 2011), low-frequency comb vibrations (Kastberger et al., 2013), or by the release of Nasonov pheromone during wave generation (Kastberger et al., 2010b).

In this study we used artificial moving stimuli that simulate aerial predators to examine the precise relationships between the visual features of the stimuli and the strength of shimmering, by varying the relative contrast and visual angles subtended by the stimuli on the bee’s eye at different ambient light levels during daylight and twilight in this facultatively nocturnal honeybee (Dyer, 1985; Somanathan et al., 2009). We predicted that the strength of shimmering will increase under the following conditions: (1) with increasing contrast between the object and the background, (2) with larger stimuli and (3) at higher ambient light levels. We also examine whether repeated exposure to potential threats (stimuli) leads to a progressive reduction in the strength of shimmering due to habituation. Since shimmering occurs in response to potential threats from predators/intruders with consequences for colony survival, we expected that habituation would not occur. We thus predicted that the shimmering response does not reduce or dampen with recurring exposure to the stimuli over trials.

## Methods

### A. dorsata colonies

The experiments were conducted in October - December 2020 and in June 2022 using two natural *A. dorsata* colonies located on the Biological Sciences building at the IISER Thiruvananthapuram campus, India (8.68°N, 77.14°E). Both colonies consisted of a single comb and nested under the roof rafters. The first colony, henceforth referred to as Hive A (1 m width, 1.2 m height), was situated 2 m above the floor level on the open terrace of the building (Fig 1A, Supp. Video) and the second colony, Hive B (1.8 m width, 0.7 m height), was 5-storeys above ground level (Fig 1B, Supp. Video). Though hives of *Apis dorsata* are often found hanging from rooftop rafters and window ledges of multi-storey buildings, accessing hives for performing controlled experiments is often risky or challenging. Both experimental hives were naturally located in their respective positions. Translocating hives to convenient locations for ease of experimentation has been done in other studies (Young et al., 2021). Since this could trigger absconding or change the baseline state of the colony, we refrained from doing so.

**Figure 1.**
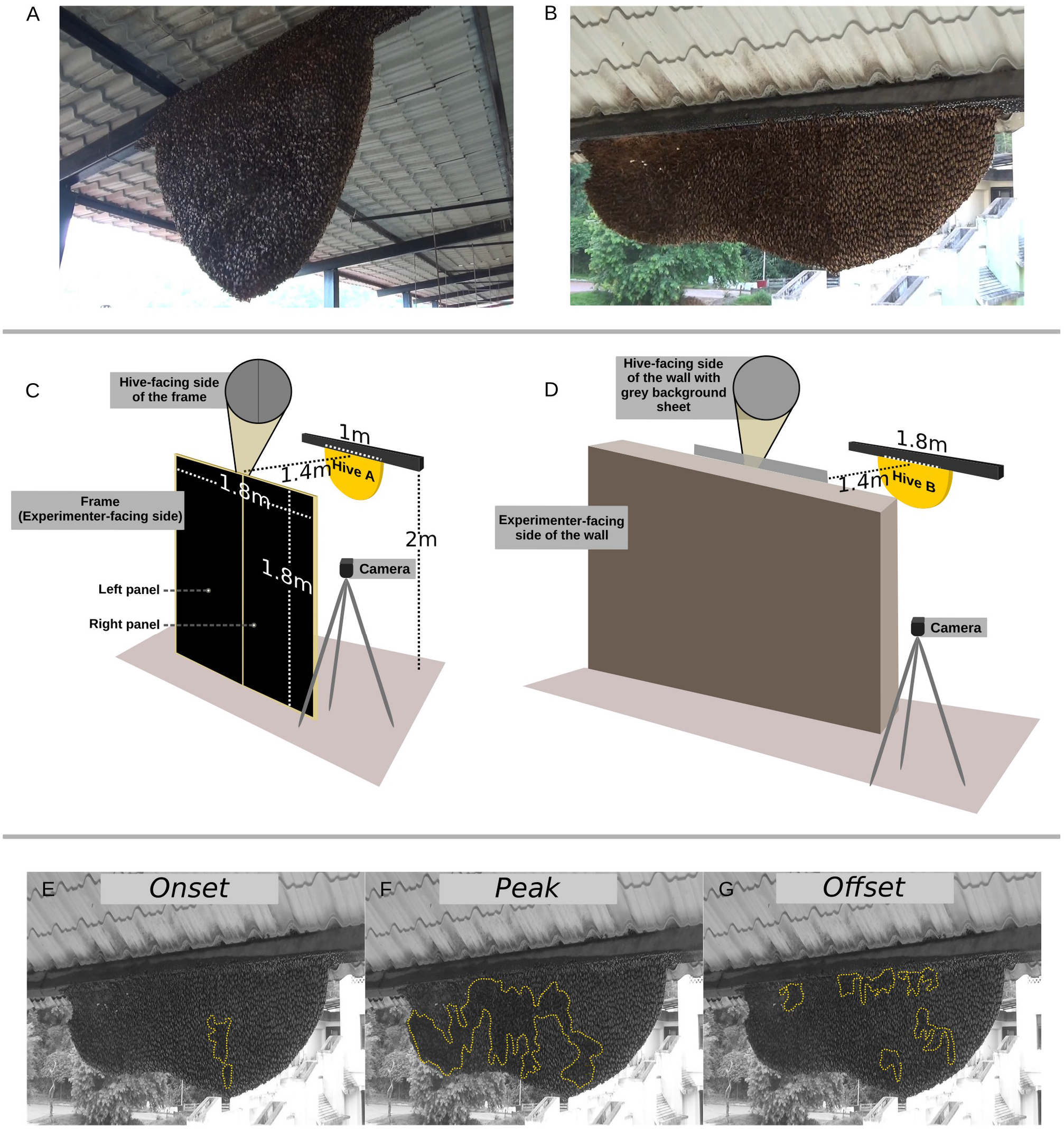
Experimental setup and quantification of shimmering response. **A-B)** Experimental colonies Hive A (A) and Hive B (B) used in the experiment. **C)** Experimental setup for Hive A. Hive-facing side of the panels were covered with the grey background, while the experimenter-facing side had a black background which could be changed by swivelling the panels. **D)** Experimental setup for Hive B. The cardboard panel covered with grey/black sheets was fastened in place from the experimenter-facing side of the wall. **E-G)** An occurrence of shimmering in Hive B. The shimmering wave (*outlined in yellow dots*) has an *onset/ascending* phase (E), *peak/climax* phase (F), and an *offset/descending* phase (G). The strength of shimmering *S*_R_ is the area covered by the climax-stage wave as a proportion of the area of the hive (F).

### Experimental setup

The stimuli used in this study were made from circular cardboard cut-outs that varied in shade (light-grey, dark-grey or black) and size (diameter: 4, 8 or 16 cm; visual angle subtended on the eye of the bee: 1.6°, 3.3° and 6.5°, respectively). The relative contrasts of the stimuli were further varied by using two different backgrounds (grey or black). In the case of Hive A, a frame that consisted of two panels that could be swivelled along their vertical axes was placed 1.4 m from the colony (Fig. 1C), and the front and rear sides of the panels were covered with grey and black papers, respectively (papers procured locally). The rear side of each stimulus was attached to a string and moved either vertically or horizontally by the experimenter standing behind the frame and out of sight of the colony. To achieve this, the string was strung around the setup vertically or horizontally. Moving the strings without a stimulus was treated as the experimental control. The position of Hive B on an overhanging roof rafter obstructed by walls and ledges prevented the use of the same arrangement. Instead, one of two backgrounds consisting of cardboard sheets (1 m x 0.6 m) covered with the black or grey papers was hung in front of the wall facing the colony during the experiments (Fig. 1D). The hive-to-background distance of 1.4 m was the same as for Hive A. The experimenter stood behind the wall and moved the stimuli horizontally with the help of a wire, and moving the wire alone acted as an experimental control. Videos of the trials were recorded using a Sony Handycam HDR CX-405.

We carried out 799 trials on Hive A, randomising the order of presentation of the stimuli across the following variables to prevent nestedness: stimulus shade, stimulus size, background colour, orientation of motion (vertical/horizontal) and side of the setup on which the trial was done (left/right). We conducted 240 trials on Hive B, randomised across stimulus shade, stimulus size, and background. Trials for both hives were carried out at least 15 minutes apart to ensure that the colonies reached a state of quiescence, and 9 – 45 trials were conducted each day between 06:00 h and 18:00 h.

The effect of ambient light intensity on shimmering was examined by moving the largest black stimulus against the grey background under dim light conditions prevailing during astronomical, nautical and civil twilight (n(Hive A) = 26, n(Hive B) = 20; illuminance between 0.002-45 lux), and the responses were compared to shimmering elicited in bright daylight (illuminance > 1000 lux). Illuminance levels were measured in lux using a Hagner Screenmaster (Hive A) and a Hagner E4-X Digital Luxmeter (Hive B).

To quantify whether habituation occurred as a result of repeated exposure within experimental days, we examined the strength of shimmering response over trials. We excluded the first data-point from each day since more than 12 hours had elapsed after the last trial.

### Relative contrasts of the stimuli against the backgrounds

The grey stimuli were made using laminated sheets of grey paper manufactured by Color-Aid Corp. The dark and light-grey stimuli were made using Color-Aid Gray-5 and Color-Aid Gray-9 respectively. Black stimuli were made from the same paper as the black background. All stimuli were pasted on to a hard cardboard base to ensure that their shapes remained unchanged over the course of the experiment. The reflectance spectra of the stimuli and backgrounds were recorded using an Ocean Insight Ocean-HDX-UV-VIS spectrophotometer connected to an Ocean Optics PX-2 pulsed Xenon light-source (Fig. S1A). The spectra were captured to a PC (Acer One 110-ICT) running the Ocean View software and saved as a spreadsheet (see electronic supplementary materials). The contrasts of the stimuli against the background were quantified using Weber contrast (O’Carroll and Wiederman, 2014)

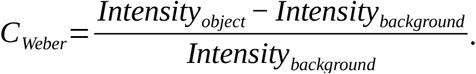

The light-grey, dark-grey and black stimuli had contrasts of 16.4, 6.6 and 0, respectively, against the black background, and 2.4, 0.5 and -0.8, respectively, against the grey background.

### Quantification of shimmering

A trial lasted 15 s and consisted of the stimulus moving 15 times from one side to the other against the front of the frame. The movement was timed using a metronome set to 60 beats per minute to maintain a constant velocity of 1 m/s (40°/s in the visual field of a bee). A shimmering response is characterised by a distinct ascending phase, a climax and a descending phase (Fig. 1E-G, (Kastberger et al., 2014)) of which only the area of the climax stage was considered for quantifying the strength of shimmering (Fig 1F). Videos were converted to AVI format using FFMPEG (at 25 fps) and analysed frame-by-frame using ImageJ software (Schneider et al., 2012). The overall strength of shimmering in a trial *S*_trial_ was quantified as

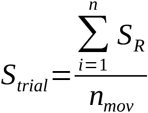

where *S*_*R*_= *Area*_*climax*_ / *Area*_*h ive*_ for the *i*_th_ climax event, and *n*_mov_ is the number of times the stimulus was moved in a trial (fixed at 15 per trial). In other words, *S*_trial_ denotes the average proportion of the hive that shimmered when the stimulus moved once.

### Statistical analyses

We compared the strength of shimmering (*S*_trial_) in relation to stimulus shade, size and background. Since the values of *S*_trial_ belong to the interval [0,1], interactions were modelled using beta regression with the hive identity as a random effect (Douma and Weedon, 2019). The side of the frame against which the stimuli were presented (left side/right side) and the direction of motion (vertical/horizontal) which were carried out only with Hive A were excluded as the shimmering response for these showed no significant differences in pairwise comparisons with Bonferroni correction (Fig. S2).

Since the beta probability distribution is defined in the open interval (0,1), and 86% of the trials elicited no shimmering (*S*_trial_=0), we rescaled *S*_trial_ in the dataset using the following equation:

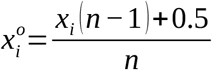

where *x*_*i*_ is the observed value, *n* is the number of trials (1039 in total) and 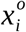 is the rescaled value for that observation. All analyses were carried out in R version 3.6.3 (R Core Team, 2021), and the beta regression analyses were done using the *glmmTMB* package (Brooks et al., 2017). We compared five beta-regression models against two null models (one with pooled data and one with hive identity as a random effect): a model with data pooled from both hives and single precision (*ϕ*) across all variables (Model 1), a model with hive identity as a fixed effect (Model 2), a hierarchical (mixed effects) model with hive identity as a random effect and single *ϕ* (Model 3), a hierarchical model with hive identity as random effect and variable *ϕ* across stimulus size (Model 4), and a hierarchical model with hive identity as random effect and variable *ϕ* across stimulus shade (Model 5). The best performing model was chosen on the basis of the Aikake Information Criterion (AIC) (see Table S1). The effect of ambient light intensity was estimated using hierarchical beta-regression modelling with light condition (daylight/twilight) being the fixed effect and hive identity as random effect (*n*_daylight_(Hive A) = 26), *n*_daylight_(Hive B) = 10, *n*_twilight_(Hive A) = 25, *n*_twilight_(Hive B) = 20), and the best performing model was chosen on the basis of AIC (see Table S2). The dataset used for the daylight condition is a subset of the data used in the previous analysis with the largest black stimulus against the grey background. Effect of habituation was analysed using linear models for both hives by estimating the impact of duration between consecutive trials on *S*_trial_.

## Results

### *A. dorsata* shimmers strongly to a dark object moving against a light background

Shimmering responses were only observed in trials where the black stimulus was moved against the grey background (Fig. 2). The hierarchical model with hive identity as random effect and variable precision (*ϕ*) across stimulus size (Model 4) was found to be the most parsimonious compared to the other models, including Model 2 where hive identity was considered as a fixed effect (Table S1, Fig. S3). Hence, we did not carry out post-hoc statistical comparisons between hives. Shimmering in Hive A was strongest with the largest black stimulus (subtending a visual angle of 6.5°, *S*_trial_ = 0.25 ± 0.12) and reduced in strength with the intermediate-sized black stimulus (3.3°, *S*_trial_ = 0.09 ± 0.10), whereas both stimuli elicited an equally strong response in the case of Hive B (*S*_trial_ (6.5°) = 0.20 ± 0.05, *S*_trial_ (3.3°) = 0.15 ± 0.04). The smallest stimulus (1.6°) did not elicit shimmering in either hive (Fig. 2). Thus, the detection threshold for a moving dark object is likely between 1.6° and 3.3°. The distinct behavioural responses and the contrasts of the stimuli (Fig. S1) suggest that bees can distinguish and respond to the stimuli on the basis of contrast and size, leading to strong shimmering responses as the visual angle subtended by the intruding object/predator increases in the visual field of the bees.

**Figure 2.**
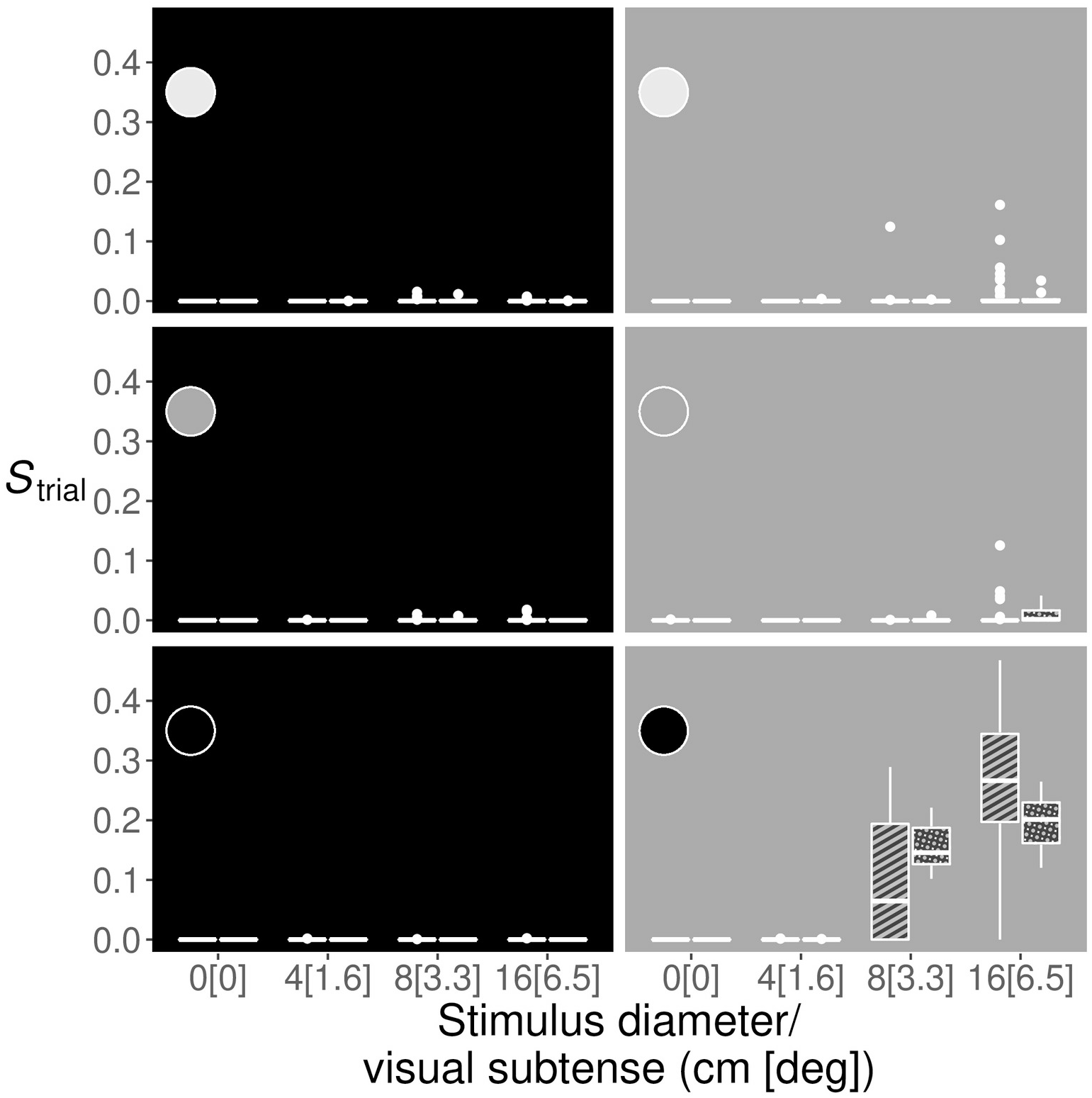
Shimmering responses in *A. dorsata* to moving stimuli against dark or light backgrounds. The relative strength of shimmering (*S*_trial_) when presented with moving circular stimuli of varying diameters/visual subtenses (x-axes) and shades (light-grey, dark-grey and black (*insets*)) against either a black (*left*) or grey (*right*) background. The *hatched boxes* correspond to Hive A, the *cross-hatched boxes* correspond to Hive B, the *hinges* (horizontal bounds of the box) correspond to the interquartile range (IQR), the *bold horizontal lines* correspond to the medians, the *whiskers* denote the upper/lower hinge ± 1.5xIQR, and the *points* outside the *whiskers* represent outliers.

### Shimmering response is less pronounced in dim-light

31 trials out of the 45 carried out during dawn and dusk with the 16 cm black stimulus against the grey background did not elicit shimmering (*n*_Hive A_ = 13 and *n*_Hive B_ = 18; illuminance between 0.1 - 3.9 lux), and the rest showed varying strengths of shimmering (illuminance between 3.9 - 45 lux). The model in which the data were pooled for both hives was the most parsimonious (Table S2). The mean strength of shimmering under bright daylight conditions was an order of magnitude higher than the response during twilight hours (*S*_trial_ (daylight) = 0.243 ± 0.101, *S*_trial_ (twilight) = 0.022 ± 0.053; Fig. 3).

**Figure 3.**
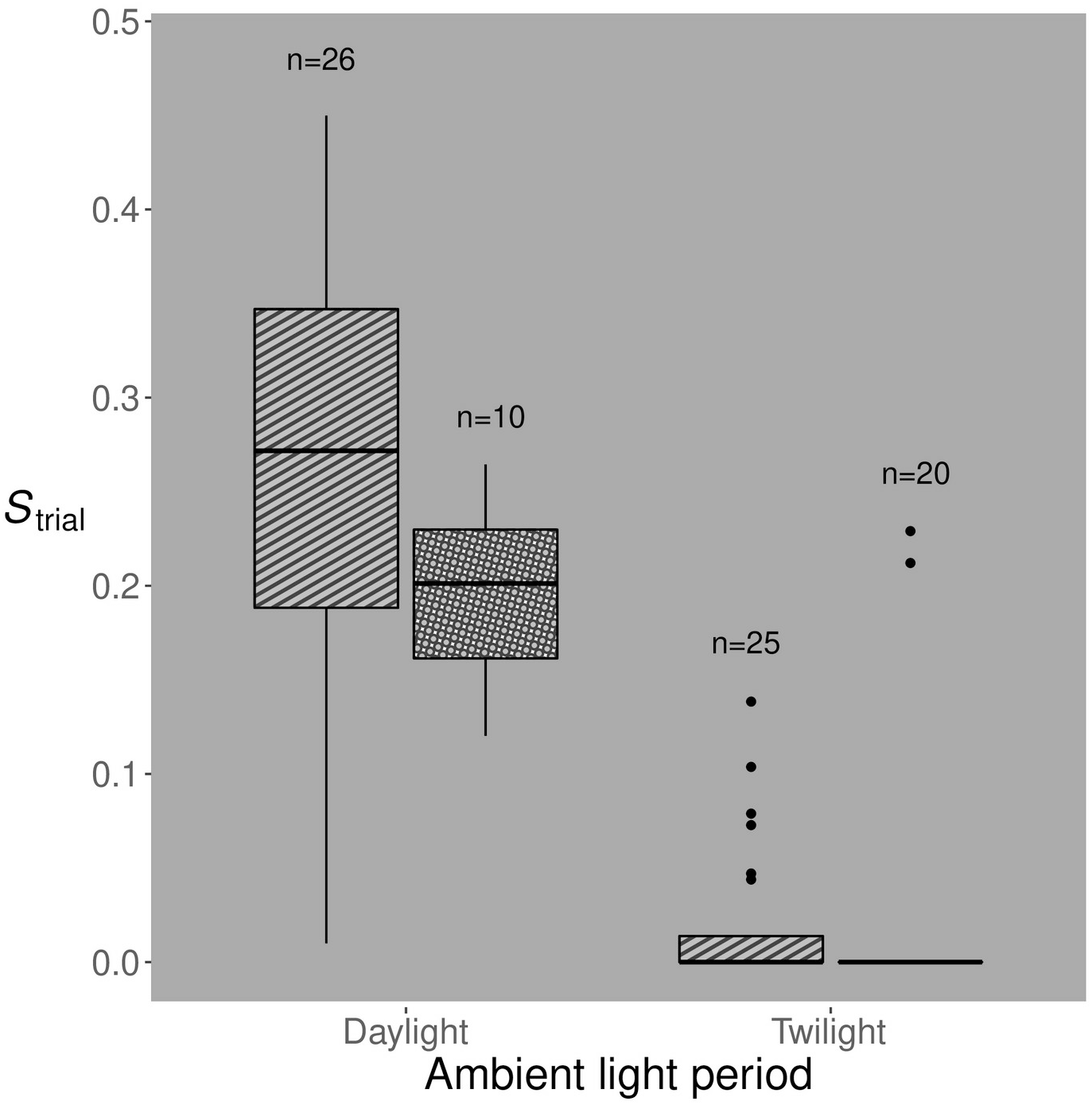
Effect of ambient light on strength of shimmering response, *S*_trial_. The colonies showed only a weak response (Hive A) and no response (Hive B) in twilight (04:45-06:45 h and 18:00-20:00 h; 0.002-45 lux) to the largest black stimulus (diameter 16 cm, subtense 6.5°) when it moved against the grey background. During daylight (07:00-17:00 h; illuminance >1000 lux), the response to the same stimulus-background combination was much more pronounced. The *hatched boxes* correspond to Hive A, the *cross-hatched boxes* correspond to Hive B, the *hinges* (horizontal bounds of the box) correspond to the interquartile range (IQR), the *bold horizontal line* corresponds to the median, the *whiskers* denote the upper/lower hinge ± 1.5xIQR, and the *points* outside the *whiskers* represent outliers.

### *A. dorsata* does not habituate with repeated exposure to the stimuli

Since *A. dorsata* shimmered only to the 8 cm and 16 cm black stimuli moving against the grey background, we excluded trials with other stimulus-background treatments from the analyses examining if habituation occurred. Since the order of presentation of stimuli and backgrounds were randomised (see Methods), the black-on-grey trials were interspersed with other treatments and the trials were often not successive. 51 trials from Hive A and 14 trials from Hive B satisfied the condition of black stimuli (8/16 cm diameter) moving against a grey background and repeating within a day (06:00 AM – 06:00 PM). We did not observe any significant change in the strength of the shimmering response with time elapsed between trials (range: 15 – 622 minutes, Fig. 4A), and there was no effect of time of the day on the strength of the shimmering response (Fig. 4B).

**Figure 4.**
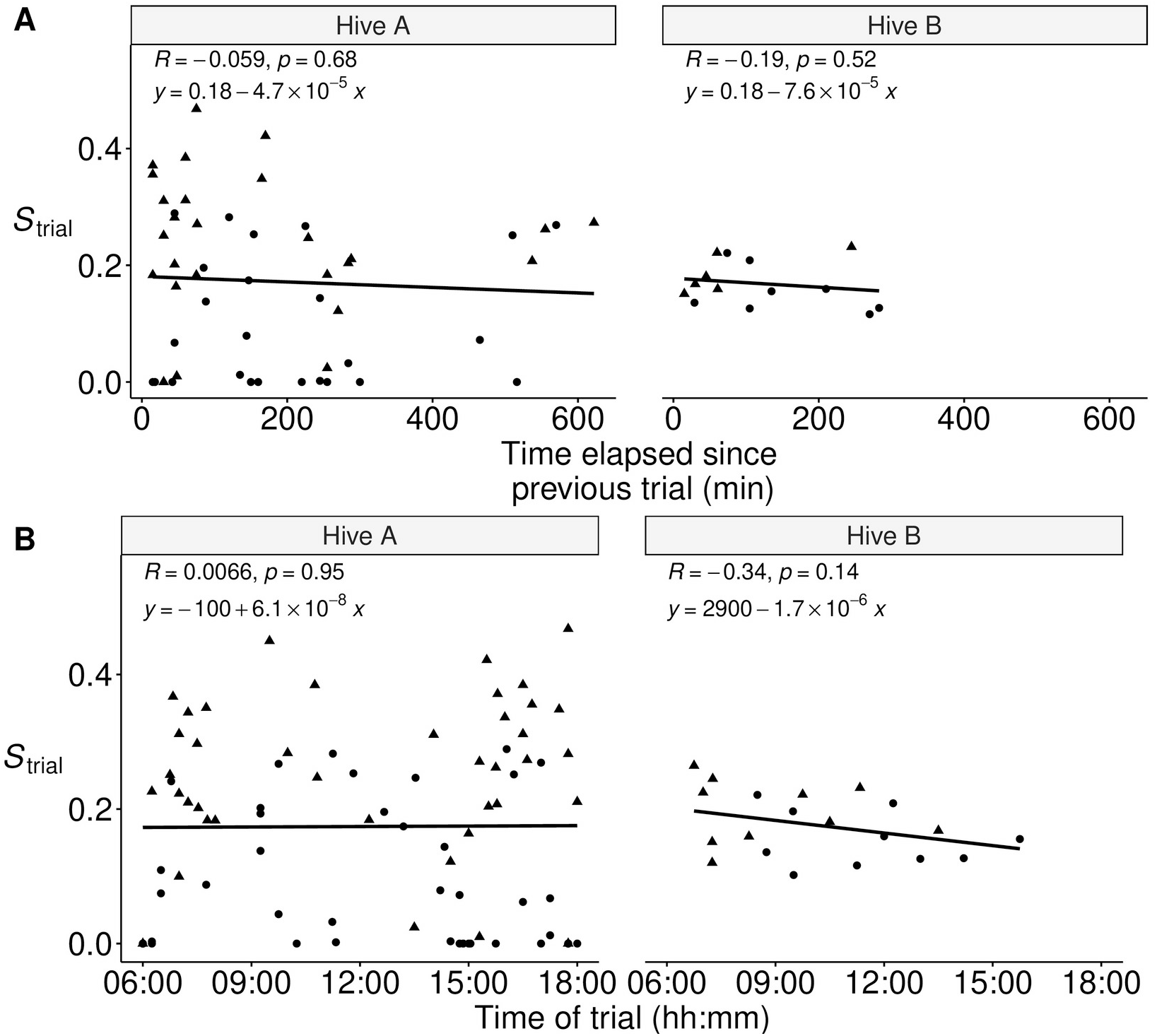
Effect of repeated exposure of stimuli on the shimmering response, *S*_trial_. **A)** Effect of time elapsed on *S*_trial_. Neither colony habituated to the black stimuli moving against the grey background. The *filled circles* (●) correspond to the 8 cm diameter black stimulus (visual subtense 3.3°), the *filled triangles* (▲) correspond to the 16 cm diameter black stimulus (visual subtense 6.5°), and the *solid line* is the linear regression of the strength of shimmering, *S*_trial_, against the time elapsed between consecutive trials. The *inset text* indicated the Pearson correlation coefficient R, the p-value and the linear relation that best explains the data. B) Effect of time of the day on *S*_trial_. Shimmering response did not exhibit any relation with the time of day when trials were conducted. The *filled circles* (●) correspond to the 8 cm diameter black stimulus (visual subtense 3.3°), the *filled triangles* (▲) correspond to the 16 cm diameter black stimulus (visual subtense 6.5°), and the *solid line* is the linear regression of the strength of shimmering, *S*_trial_, against the time of the day. The *inset text* indicated the Pearson correlation coefficient R, the p-value and the linear relation that best explains the data.

## Discussion

Defence behaviour in honeybees involves several sensory modalities (Fuchs and Tautz, 2011), and shimmering is predominantly mediated by visual cues (Kastberger et al., 2010a; Kastberger et al., 2011). When we simulated predator movement using artificial stimuli, the strongest shimmering responses were obtained only in bright ambient light when the stimulus was much darker than the background. This is unlike the role of contrast in the context of foraging (in *A. mellifera*), where the stimulus can be either brighter or darker than the background and yet elicit similar responses (de Ibarra et al., 2000). We also found that the strength of shimmering decreased as stimulus size reduced and resulted in no response with the smallest stimulus. Repeated exposure to the experimental stimuli did not lead to a change in the strength of the shimmering response in *A. dorsata*, although habituation or sensitisation occurs in some invertebrates in response to repeated exposure to a threat (Evans et al., 2019).

Detection of dark objects moving against bright backgrounds (e.g. the sky) has been reported in several insects, including bees, although mostly in the context of mate-detection or predation (Bergman et al., 2015; Sauseng et al., 2003). Male carpenter bees (*Xylocopa tenuiscapa*) detect and chase inanimate projectiles, mistaking them to be potential mates or rivals, when these objects subtend very small visual angles of 0.4° - 1° (which darken the visual field of a single ommatidium by as little as 2%) (Somanathan et al., 2017). *A. mellifera* drones fly towards moving objects subtending as little as 0.4° (which darkens the visual field by just 8%) (Vallet and Coles, 1993). *A. mellifera* workers can also detect a small black stimulus (0.6° x0.6°) moving at 65°/s against a white background (Rigosi et al., 2017), which is significantly smaller than their detection threshold during foraging (3.7°-5°) (Giurfa and Vorobyev, 1998). Anatomical estimates from the compound eyes of *A. dorsata* suggest coarser visual resolution compared to other honeybees (Somanathan et al., 2009), which agrees with recent estimates of detection thresholds in *A. dorsata* in the context of foraging (achromatic: 5.7°, chromatic: 12.4°, Balamurali et al. *in prep*).

Our experiments suggest that *A. dorsata* shimmers in response to dark objects moving against a bright background that subtend a visual angle of at least 3.3° (and likely even smaller, between 1.6° and 3.3°), which is smaller than their achromatic detection threshold while foraging. Using a conservative detection threshold of 3.3° for shimmering behaviour, and assuming that a colony shimmers to a hornet (a common predator) which is ∼2.3-2.7 cm in length (Girish and Srinivasan, 2010), the detection-distance threshold corresponds to ∼40-47 cm, which agrees with results from real-world interactions involving free-flying hornets near *A. dorsata* colonies (Kastberger et al., 2008). Besides predators, *A. dorsata* also shimmer in response to intruders such as non-nestmate bees who display erratic flight patterns and splay their legs outwards while doing so (Weihmann et al., 2014). Based on a length of ∼2 cm for an *A. dorsata* worker (pers. obs.) and a detection threshld of 3.3°, we speculate that the distance at which shimmering is triggered is around 35 cm. This suggests that shimmering could be a general response to aerial threats.

Open-nesting honeybees such as *A. dorsata* are likely to perceive flying predators as dark objects moving against the bright sky. Hornets are major predators of *A. dorsata* hives (Kastberger et al., 2013; Seeley et al., 1982), and *Vespa tropica* (the greater banded hornet) is a common predator in our study location (Balamurali et al., 2021). Moreover, hornets can act as strong evolutionary drivers in the evolution of specialised anti-predatory responses in honeybees (Cappa et al., 2021; Ken et al., 2005; Mattila et al., 2020; Mattila et al., 2021; Papachristoforou et al., 2007; Tan et al., 2013). Shimmering resembles the anti-hornet ‘I-see-you’ display found in the sympatric cavity-nesting *A. cerana* in which guard bees at the entrance shake their abdomens in a to-and-fro motion in the presence of hornets, which supposedly warns the predator that the bees have detected its presence (Tan et al., 2013). Simulating this threat with black discs resulted in strong abdomen-shaking responses in *A. cerana* to stimuli of size 8° moving at 140°/s (Koeniger and Fuchs, 1973, as cited in Fuchs and Tautz, 2011)), which agrees with the achromatic visual detection threshold of 7.7° in *A. cerana* (Meena et al., 2021). In the context of an anti-predatory response such as shimmering, habituation may prove costly, and we found that the shimmering response does not reduce in strength after repeated exposure to the stimuli, which is advantageous to ward off repeated approaches/attacks by predators.

Although in our study we found that the shimmering response increased with larger stimulus size, up to a maximum target size of 6.5° used in our experiments, we cannot rule out the possibility that the response increases further for even larger targets, or simply saturates. Beyond a certain size threshold (which is between 6.5° (current study) and 9.5° (Koeniger et al., 2017)), the colony responds to the object with stinging attacks (Koeniger et al., 2017), which resembles the defensive response of *A. dorsata* against much larger predators such as birds (Kastberger and Sharma, 2000). The findings from our study using moving objects are consistent with the notion that the shimmering behaviour of the giant honeybee is a response to an aerial predator moving in front of the hive against a bright sky background.

## Supporting information

Datasets of shimmering response and reflectance spectra of stimuli and backgrounds

Supplementary methods and results

Video of the shimmering response in both hives used in this study.

Contains the code for all analyses and plots in the main text and the supplementary materials

## Acknowledgements

SV was funded by the Council for Scientific and Industrial Research, India, SV and HS were funded by IISER Thiruvananthapuram and EJW was funded by the Swedish Research Council (2016-04014). We thank GS Balamurali for his valuable inputs, and PC Anand for helping with the pilot experiments for this study. We also thank three anonymous reviewers for their valuable comments on the manuscript.

