## Supplementary methods and results for "Defensive shimmering responses in *Apis dorsata* are triggered by dark stimuli moving against a bright background"

#### Reflectance spectra and contrasts of stimuli

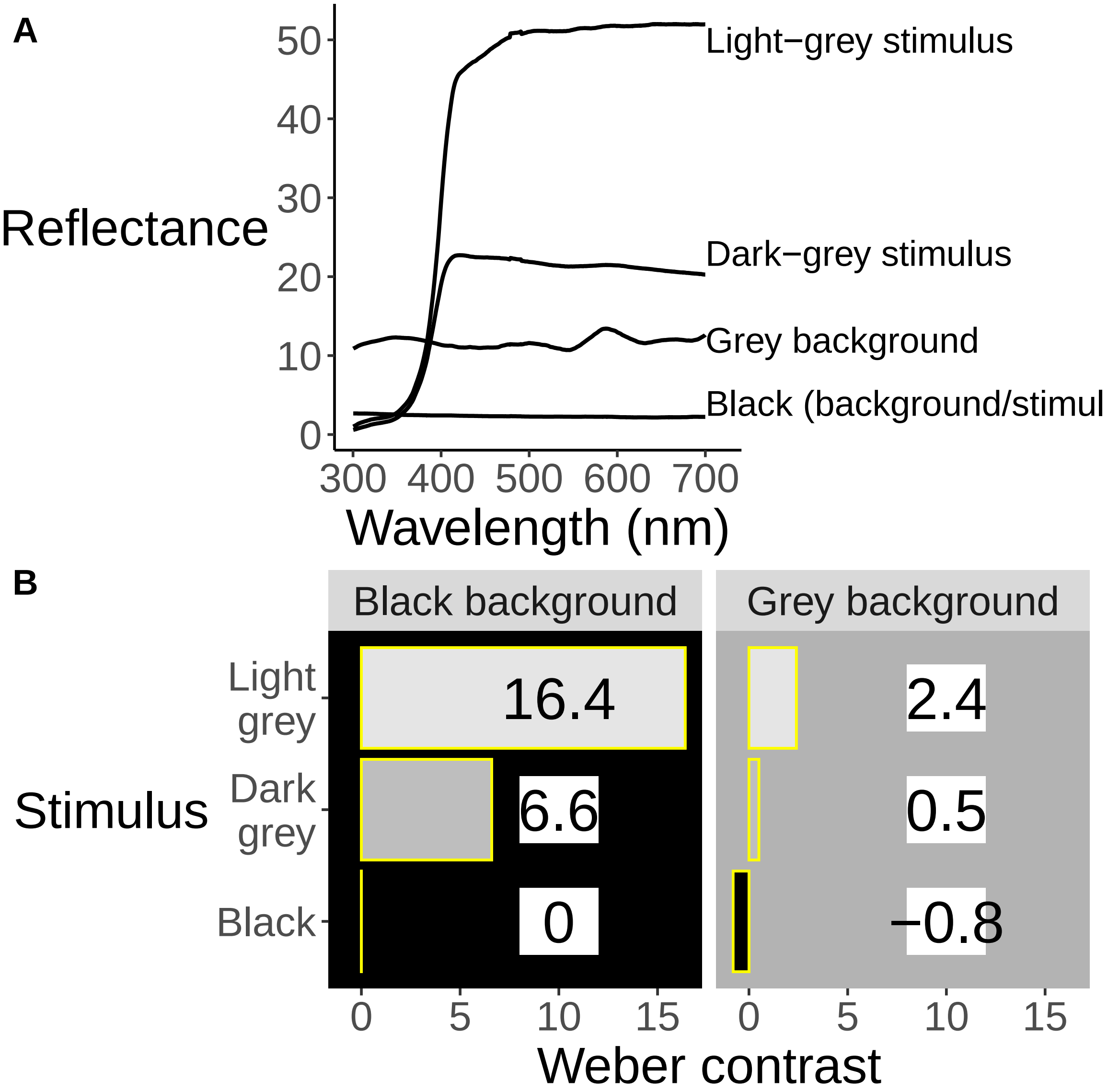


**Figure S1. A)** The reflectance spectra of the stimuli and backgrounds were recorded using an Ocean Insight

Ocean-HDX-UV-VIS spectrophotometer connected to an Ocean Optics PX-2 pulsed Xenon light-source. The spectra were captured to a PC (Acer One 110-ICT) running the Ocean View software and saved as a spreadsheet. **B)** The Weber contrasts of the stimuli against the grey or black backgrounds were calculated using the formula provided in the main text, adapted from O'Carroll & Wiederman 2014. The contrast values are written inside/beside the corresponding *bars*.

### Supplementary results:

#### Effect of orientation of motion and side of the trial on shimmering response

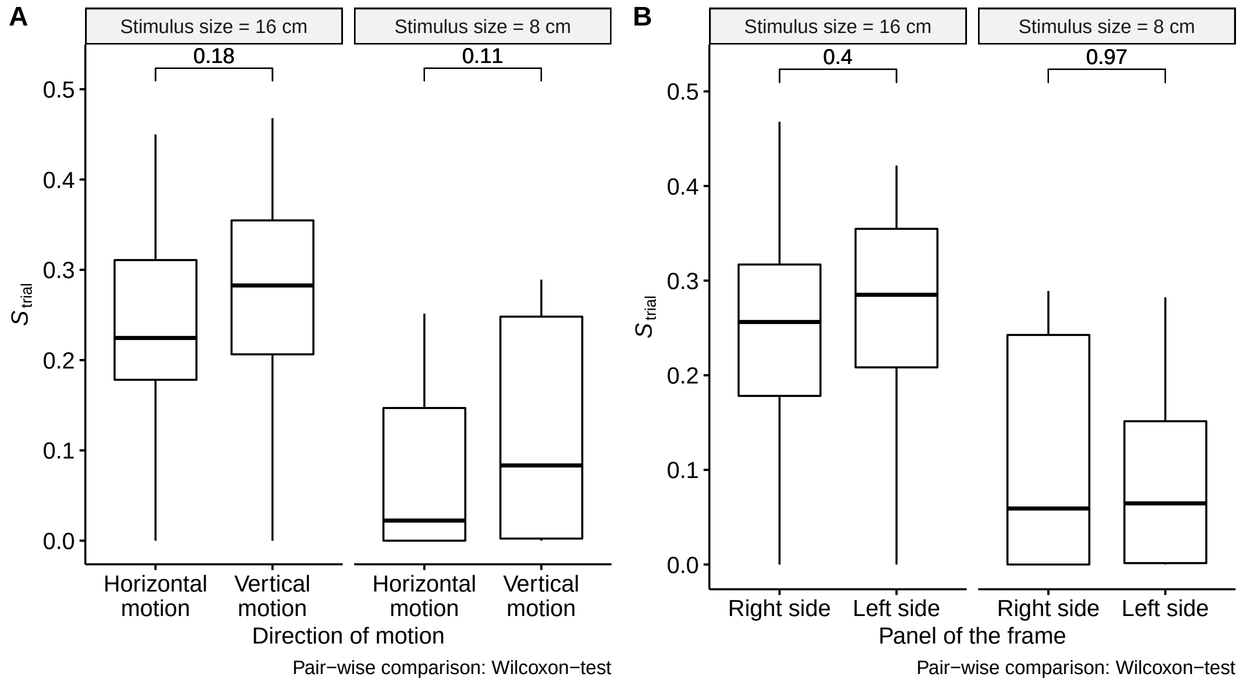


**Figure S2.** Pairwise comparison of shimmering responses between direction of motion **(A)** and side of the panel **(B)**. $S_{\text{trial}}$ refers to the shimmering strength of each trial (refer to main text). The *numbers in boldface* correspond to the p-value of the comparison. There was no effect of direction of motion or side of panel for all stimuli sizes. Hence these two variables were excluded from the final model.

### Beta-regression modelling and comparison between models

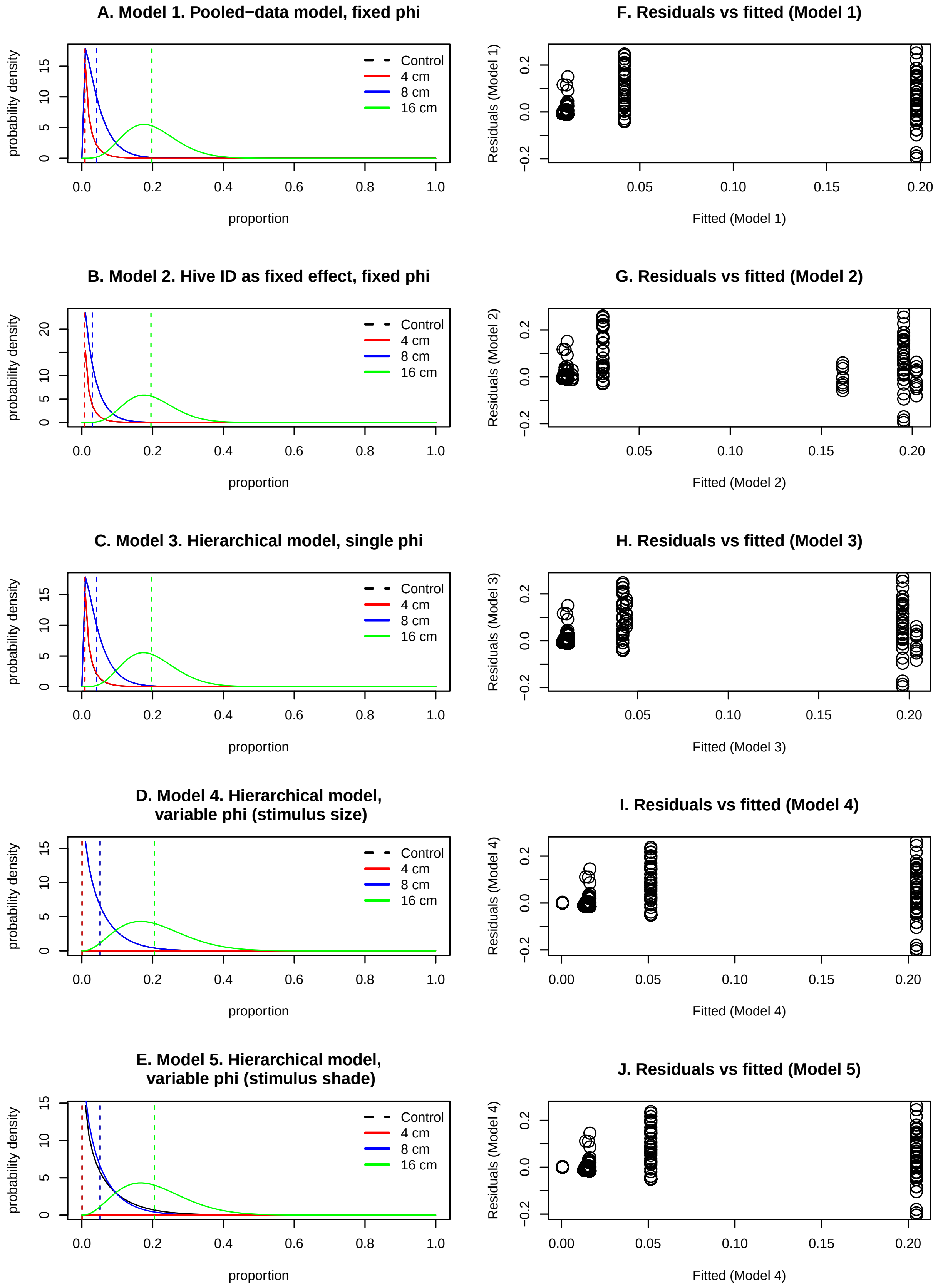

**Figure S3. A-E) Probability densities of shimmering response to black stimuli against a grey background predicted from the MLE parameters for Models 1-5.** *'Control'* denotes the experimental condition where the string/wire alone was moved, and 4 cm, 8 cm and 16 cm are the diameters of the circular black stimuli. The *coloured lines* are density distributions of the shimmering response back-transformed for the sake of comparison with the original data. **(F-J) Standardised residuals against the fitted values** The spread of residuals is plotted along the y-axis against the experimental treatments on the x-axis.

| | dAIC | Degrees of freedom | Model description |
|---|---|---|---|
| glm.4 | 0.000 | 29 | Hierarchical, variable precision (stimulus size) |
| glm.5 | 1584.676 | 28 | Hierarchical, variable precision (stimulus shade) |
| glm.2 | 2124.989 | 49 | Hive identity as fixed effect |
| glm.1 | 2136.250 | 25 | Pooled-data, single precision |
| glm.3 | 2138.109 | 26 | Hierarchical, single precision |
| glm.0 | 2591.786 | 2 | Pooled-data null model |
| glm.01 | 2593.786 | 3 | Hierarchical null model |

**Table S1. Comparison of the beta-regression models quantifying the shimmering response of black stimuli against grey background during bright light conditions**

| | dAIC | Degrees of freedom | Model description |
|---|---|---|---|
| glm.6 | 0.00000 | 3 | Pooled-data model |
| glm.7 | 2.00000 | 4 | Hierarchical model |
| glm.02 | 88.15653 | 2 | Pooled-data null model |
| glm.03 | 89.86991 | 3 | Hierarchical null model |

**Table S2. Beta-regression models comparing the shimmering response during daylight to that occurring in twilight**

*This document was generated from the supplementary markdown file. The details of the session and the packages used to generate this file are printed below*

***Session and packages info***

```
##  setting  value
##  version  R version 4.1.3 (2022-03-10)
##  os       Fedora Linux 35 (Workstation Edition)
##  system   x86_64, linux-gnu
##  ui       X11
##  language en_GB
##  collate  en_IN.UTF-8
##  ctype    en_IN.UTF-8
##  tz       Asia/Kolkata
##  date     2022-06-23
##  pandoc   2.14.0.3 @ /usr/libexec/rstudio/bin/pandoc/ (via rmarkdown)
```

```
##              package ondiskversion
## bbmle          bbmle        1.0.25
## boot            boot        1.3.28
## cowplot      cowplot         1.1.1
## dplyr          dplyr         1.0.7
## emmeans      emmeans       1.7.1.1
## EnvStats    EnvStats         2.7.0
## ggpattern  ggpattern       0.4.3.3
## ggplot2      ggplot2         3.3.6
## ggpubr        ggpubr         0.4.0
## glmmTMB      glmmTMB       1.1.2.3
## latex2exp  latex2exp         0.5.0
## lsmeans      lsmeans        2.30.0
## lubridate  lubridate         1.8.0
## stringr      stringr         1.4.0
## tidyr          tidyr         1.1.4
##                                                                     source
## bbmle                                                       CRAN (R 4.1.3)
## boot                                                        CRAN (R 4.1.3)
## cowplot                                                     CRAN (R 4.1.2)
## dplyr                                                       CRAN (R 4.1.1)
## emmeans                                                     CRAN (R 4.1.2)
## EnvStats                                                    CRAN (R 4.1.3)
## ggpattern Github (coolbutuseless/ggpattern@1f46c8bc0c547cdbc3cc051e81e94625a1e0f1a6)
## ggplot2                                                     CRAN (R 4.1.3)
## ggpubr                                                      CRAN (R 4.1.2)
## glmmTMB                                                     CRAN (R 4.1.2)
## latex2exp                                                   CRAN (R 4.1.2)
## lsmeans                                                     CRAN (R 4.1.3)
## lubridate                                                   CRAN (R 4.1.2)
## stringr                                                     CRAN (R 4.1.2)
## tidyr                                                       CRAN (R 4.1.1)
```

*The last section of the accompanying markdown file* (sajesh_etal_2022_ESM.Rmd) *contains the code for Figs. 2-4 in the main text. The plots can be generated by changing the value of the* `include` *argument in the corresponding code-chunk to* `TRUE`
