## Supplementary material for "Defensive shimmering responses in *Apis dorsata* are triggered by dark stimuli moving against a bright background": Contains the code for all analyses and plots in the main text and the supplementary materials: sajesh_etal_2022_rev_ESM.Rmd

````
---
title: "Supplementary Materials"
author: "Sajesh Vijayan, Eric J. Warrant & Hema Somanathan"
date: ''
output:
  pdf_document: default
  word_document: default
  html_document: default
subtitle: 'Defensive shimmering responses in *Apis dorsata* are triggered by dark stimuli moving against a bright background'
---

```{r setup, include=FALSE}
rm(list = ls())

knitr::opts_chunk$set(echo = TRUE)
library(dplyr)
library(tidyr)
library(stringr)
library(lubridate)
library(lsmeans)
library(glmmTMB)
library(boot)
library(ggplot2)
library(ggpubr)
library(latex2exp)
library(ggpubr)
library(cowplot)
library(bbmle)
library(ggpattern)
library(EnvStats)


# loading the datasets
refl_spectra <- read.csv("./sajesh_etal_2022_spectra.csv") # col1: WL, col2: black, col4: dark-grey, col5: light-grey
shimData <- read.csv("./sajesh_etal_2022_shimmerData.csv", colClasses = c(cardSize = "factor"))

lightData <- read.csv("./sajesh_etal_2022_shimmerLowLight.csv")

# function to transform data for beta-regression modelling
transform01 <- function(x) {
  (x * (length(x) - 1) + 0.5) / (length(x))
}

# functions to help visualise beta-regression models
dbeta2 <- function(X, mu, phi, ...) {
  dbeta(X, shape1 = mu * phi, shape2 = (1 - mu) * phi, ...)
}

rbeta2 <- function(N, mu, phi, ...) {
  rbeta(N, shape1 = mu * phi, shape2 = (1 - mu) * phi, ...)
}
```


## Supplementary methods

### Reflectance spectra and contrasts of stimuli

```{r reflSpectra, echo=F, message=FALSE, warning=FALSE, fig.height=6, fig.width=6}
reflTidyWide <- refl_spectra %>%
  select(wavelength1, starts_with("reflectance"))

names(reflTidyWide) <- c("wavelength", "black", "greyBG", "dark-grey", "grey")

### Plotting reflectance
reflPlot <- reflTidyWide %>%
  gather(cardID, reflectance, -wavelength) %>%
  filter(wavelength <= 700 & wavelength >=300) %>%
  ggplot(aes(wavelength, reflectance, colour = cardID)) +
  geom_path(size=0.75)+
  scale_color_manual(name = "Card colour",
                     values = c("black" = "black",
                                "greyBG" = "black",
                                "dark-grey" = "black",
                                "grey" = "black"))+
  scale_x_continuous(limits = c(300,720)) +
  labs(x = "Wavelength (nm)", y = "Reflectance")+
  coord_cartesian(xlim = c(300,720), # This focuses the x-axis on the range of interest
                  clip = 'off') +   # This keeps the labels from disappearing
  annotate("text", x = 700, y = c(4,12,23,50),
           size = 5,
           label = c("Black (background/stimulus)",
                     "Grey background","Dark-grey stimulus",
                     "Light-grey stimulus"),
           hjust = 0)+
  theme(
    text = element_text(size=20),
    panel.background = element_rect(fill = "transparent"), # bg of the panel
    plot.background = element_rect(fill = "transparent", color = NA), # bg of the plot
    panel.grid.major = element_blank(), # get rid of major grid
    panel.grid.minor = element_blank(), # get rid of minor grid
    legend.background = element_rect(fill = "transparent"), # get rid of legend bg
    legend.box.background = element_rect(colour = NA,
                                         fill = "transparent"), # get rid of legend panel bg
    axis.title.y=element_text(angle=0, vjust = 0.5),
    plot.margin = unit(c(0,5,0,0), "cm"),
    axis.line.x = element_line(size = 0.5, linetype = "solid", colour = "black"),
    axis.line.y = element_line(size = 0.5, linetype = "solid", colour = "black"),
    legend.position = "none")

### Calculating Weber contrasts
intensityDF <- reflTidyWide %>%
  filter(wavelength >= 300 & wavelength <=700) %>%
  select(-wavelength) %>%
  colSums() %>%
  as_tibble() %>%
  mutate(colour  = c("black", "gBG", "DG", "LG")) %>%
  mutate(sumIntensity = value) %>%
  select(colour, sumIntensity) %>%
  as.data.frame()

webCon = data.frame(stimCol = NA, bgCol = NA, stimIntensity = NA , weberCon = NA)
for(i in c("black", "DG", "LG")){
  for(j in c("black", "gBG")){
    stimCol = i
    bgCol = j
    stimIntensity <- intensityDF$sumIntensity[intensityDF$colour == i]
    bgIntensity <- intensityDF$sumIntensity[intensityDF$colour == j]
    weberCon = (stimIntensity - bgIntensity)/bgIntensity
    newEntry  = c(stimCol, bgCol, stimIntensity, weberCon)
    webCon = rbind(webCon, newEntry)
  }
}

### Plotting the contrasts
webCon <- webCon %>%
  drop_na() %>%
  group_by(bgCol) %>%
  arrange(bgCol) %>%
  mutate(weberCon=round(as.numeric(weberCon),1))
  
contrastPlots <- ggplot(webCon, aes(as.numeric(weberCon), stimCol, fill = stimCol))+
  geom_rect(data = subset(webCon, bgCol == "black"),
            xmin = -Inf,xmax = Inf,
            ymin = -Inf,ymax = Inf, fill = "black") +
  geom_rect(data = subset(webCon, bgCol == "gBG"),
            xmin = -Inf,xmax = Inf,
            ymin = -Inf,ymax = Inf, fill = "gray70") +
  geom_rect(xmin = 8,xmax = 12,
            ymin = 0.7,ymax = 1.3, fill = "white") +
  geom_rect(xmin = 8,xmax = 12,
            ymin = 1.7,ymax = 2.3, fill = "white") +
  geom_rect(xmin = 8,xmax = 12,
            ymin = 2.7,ymax = 3.3, fill = "white") +
  geom_bar(color = "yellow", stat = "identity")+geom_bar(color = "yellow", stat = "identity")+
  scale_y_discrete(labels=c("LG" = "Light\n grey",
                            "DG" = "Dark\n grey",
                            "black" = "Black"))+
  scale_fill_manual(values = c("LG" = "gray90",
                               "DG" = "grey",
                               "black" = "black")) +
  labs(x = "Weber contrast", y="Stimulus") +
  facet_grid(~bgCol, 
             labeller = labeller(bgCol = c("black"="Black background",
                                               "gBG"="Grey background"))) +
  theme(
    text = element_text(size=20),
    panel.background = element_rect(fill = "transparent"), # bg of the panel
    plot.background = element_rect(fill = "transparent", color = NA), # bg of the plot
    panel.grid.major = element_blank(), # get rid of major grid
    panel.grid.minor = element_blank(), # get rid of minor grid
    legend.background = element_rect(fill = "transparent"), # get rid of legend bg
    legend.box.background = element_rect(colour = NA,
                                         fill = "transparent"), # get rid of legend panel bg
    axis.title.y=element_text(angle=0, vjust = 0.5),
    legend.position = "none"
  ) +
  geom_text(data = webCon, aes(x = 10), label=webCon$weberCon, size = 8, color = "black")


plot_grid(reflPlot, contrastPlots, labels = c('A','B'), ncol = 1)
```

<br>
**Figure S1. A)** The reflectance spectra of the stimuli and backgrounds were recorded using an Ocean Insight Ocean-HDX-UV-VIS spectrophotometer connected to an Ocean Optics PX-2 pulsed Xenon light-source. The spectra were captured to a PC (Acer One 110-ICT) running the Ocean View software and saved as a [spreadsheet](./sajesh_etal_2022_spectra.csv).  **B)** The Weber contrasts of the stimuli against the grey or black backgrounds were calculated using the formula provided in the main text, adapted from [O'Carroll & Wiederman 2014](http://doi.org/10.1098/rstb.2013.0043). The contrast values are written inside/beside the corresponding *bars*.

<br>
<br>
<br>

## Supplementary results:
### Effect of orientation of motion and side of the trial on shimmering response
```{r directionAndSection, fig.width=9, fig.height=5,echo=FALSE, message=FALSE, warning=FALSE}
dirPlot <- shimData %>%
  filter(colourCards == "B" & bgColour == "Gray" & (cardSize == "8" | cardSize == "16") & hiveID == "A") %>%
  ggboxplot(x = "moveDir", y = "avgArea")+
  scale_x_discrete(labels = c("H" = "Horizontal\nmotion",
                              "V" = "Vertical\nmotion"))+
  facet_grid(~cardSize, labeller = as_labeller(c("8" = "Stimulus size = 8 cm",
                                                         "16" = "Stimulus size = 16 cm"))) +
  stat_compare_means(comparisons = list(c("H","V"), c("H","V"),c("H","V"),c("H","V")), 
                     label.y = 0.5, method = "wilcox", p.adjust.methods = "bonferroni")+
  theme(text = element_text(size = 12),
        legend.position = "None")+
  labs(x = "Direction of motion", y = TeX('$\\textit{S}_{trial}$'), caption = "Pair-wise comparison: Wilcoxon-test")

panelPlot <- shimData %>%
  filter(colourCards == "B" & bgColour == "Gray" & (cardSize == "8" | cardSize == "16"),
         hiveID == "A") %>%
  ggboxplot(x = "moveSec", y = "avgArea")+
  scale_x_discrete(labels = c("L" = "Left side",
                              "R" = "Right side"))+
  facet_grid(~cardSize, labeller = as_labeller(c("8" = "Stimulus size = 8 cm",
                                                 "16" = "Stimulus size = 16 cm"))) +
  stat_compare_means(comparisons = list(c("L","R"), c("L","R"),c("L","R"),c("L","R")), 
                     label.y = 0.5, method = "wilcox")+
  theme(text = element_text(size = 12),
        legend.position = "None")+
  labs(x = "Panel of the frame", y = TeX('$\\textit{S}_{trial}$'), caption = "Pair-wise comparison: Wilcoxon-test")

plot_grid(dirPlot, panelPlot, labels = c('A','B'))
```

**Figure S2.** Pairwise comparison of shimmering responses between direction of motion **(A)** and side of the panel **(B)**. *S*~trial~ refers to the shimmering strength of each trial (refer to main text). The *numbers in boldface* correspond to the p-value of the comparison. There was no effect of direction of motion or side of panel for all stimuli sizes. Hence these two variables were excluded from the final model.


### Beta-regression modelling and comparison between models

```{r modelsToCompare, echo=FALSE, message=FALSE, warning=FALSE}

##### Effect of relative contrast and size of stimuli #####
# Data cleaning
shimData$trialTime <- as.POSIXct(shimData$trialTime, format = "%Y-%m-%d %H:%M:%S")
shimData$cardSize <- factor(shimData$cardSize, levels = c("0", "4", "8", "16"))

# reate a new vector with a random distribution of the three non-Control colour card values
set.seed(5)
newCon <- sample(c("B", "G1", "G3"), 80, replace = T)

# replace the existing values with the newly created vector
shimData$colourCards[shimData$colourCards == "Control"] = newCon
shimData$colourCards <- as.factor(as.character(shimData$colourCards))
shimData$colourCards <- factor(shimData$colourCards, 
                               levels = c("G1", "G3", "B"))

shimData$trialDate <- as.Date(shimData$trialTime, tz = "Asia/Kolkata")
shimData$trialTimes <- lubridate::ymd_hms(shimData$trialTime) %>% 
  format(format = "%H:%M:%S") %>% 
  as.POSIXct(format = "%H:%M:%S")

# scaling avgArea
shimData$avgArea.SCALED <- transform01(shimData$avgArea)


# Beta-regression null model
glm.0 <- glmmTMB(avgArea.SCALED ~ 1, data = shimData, 
                 family = list(family = "beta", link = "logit"))


# Beta-regression null model with hiveID as random effect
glm.01 <- glmmTMB(avgArea.SCALED ~ 1 + (1 | hiveID), data = shimData, 
                 family = list(family = "beta", link = "logit"))

# Beta-regression with single precision
glm.1 <- glmmTMB(avgArea.SCALED ~ colourCards*cardSize*bgColour, data = shimData, 
                 family = list(family = "beta", link = "logit"))

# Beta-regression with hive ID (hiveID) as fixed effect
glm.2 <- glmmTMB(avgArea.SCALED ~ colourCards*cardSize*bgColour*hiveID, 
                 data = shimData, 
                 family = list(family = "beta", link = "logit"))

# Beta-regression with hive ID (hiveID) as random effect
glm.3 <- glmmTMB(avgArea.SCALED ~ colourCards*cardSize*bgColour + (1 | hiveID), 
                 data = shimData, 
                 family = list(family = "beta", link = "logit"))

# Beta-regression with random effect and precision varying along stimulus size (card size)
glm.4 <- update(glm.3, dispformula = ~ cardSize)

# Beta-regression with random effect and precision varying along stimulus shade (colourCards)
glm.5 <- update(glm.3, dispformula = ~ colourCards)


##### Effect of ambient light on shimmering #####
brightDataA<-shimData %>%
  filter(colourCards == "B" &
           cardSize == "16" &
           bgColour == "Gray" &
           hiveID == "A") %>%
  mutate(since_midnight = hour(trialTime) * 60 + minute(trialTime)) %>%
  filter(since_midnight >= (7*60+30) & since_midnight < 17*60) %>% 
  mutate(lightCond = "daylight") %>%
  select(avgArea, lightCond, hiveID)

brightDataB <- shimData %>%
  filter(colourCards == "B" &
           cardSize == "16" &
           bgColour == "Gray" &
           hiveID == "B") %>%
  mutate(lightCond = "daylight") %>%
  select(avgArea, lightCond, hiveID)

dimData <- lightData %>%
  na.omit() %>%
  mutate(lightCond = "twilight") %>%
  select(avgArea, lightCond, hiveID)

ambLightData <- rbind(brightDataA, brightDataB, dimData)

ambLightData[ambLightData == "daylight"] <- "Daylight"
ambLightData[ambLightData == "twilight"] <- "Twilight"

ambLightData$avgArea.SCALED <- transform01(ambLightData$avgArea)


# Beta-regression with single precision
glm.02 <- glmmTMB(avgArea.SCALED ~ 1, data = ambLightData,
                 family = list(family = "beta", link = "logit"))
# Beta-regression with single precision
glm.03 <- glmmTMB(avgArea.SCALED ~ 1 + (1 | hiveID), data = ambLightData,
                 family = list(family = "beta", link = "logit"))

# Beta-regression with single precision
glm.6 <- glmmTMB(avgArea.SCALED ~ lightCond, data = ambLightData,
                 family = list(family = "beta", link = "logit"))

# Beta-regression with hive ID (hiveID) as random effect
glm.7 <- glmmTMB(avgArea.SCALED ~ lightCond + (1 | hiveID), 
                 data = ambLightData, family = list(family = "beta", link = "logit"))

```


```{r betaPlots, echo=FALSE, message=FALSE, warning=FALSE, error=TRUE, fig.height=11, fig.width=8}
coef_to_mean <- function(model) {
  vals <- fixef(model)$cond
  out <- c(vals[1], vals[2:length(vals)] + vals[1])
  return(out)
}

old <- par(mfrow = c(5, 2))

##### Model 1 distribution #####
muphi.betaModel1 <- unique(data.frame(
  mu = fitted(glm.1),
  phi = predict(glm.1, type = "disp"),
  cardSize = shimData$cardSize,
  colourCards = shimData$colourCards,
  bgColour = shimData$bgColour
))

muphi.betaModel1.SUB <- muphi.betaModel1[muphi.betaModel1$colourCards == "B" & muphi.betaModel1$bgColour == "Gray",]


curve(dbeta2(x, muphi.betaModel1.SUB[1, 1], muphi.betaModel1.SUB[1, 2]),
      xlab = "proportion", ylab = "probability density",
      main = "A. Model 1. Pooled-data model, fixed phi"
)

abline(v = muphi.betaModel1.SUB[1, 1], lty = 2)

for (i in 1:4) {
  curve(dbeta2(x, muphi.betaModel1.SUB[i, 1], muphi.betaModel1.SUB[i, 2]), add = T, 
        col = c("blue", "black", "red", "green")[i], 
        lty = 1)
  abline(v = muphi.betaModel1.SUB[i, 1], col = c("blue", "black", "red", "green")[i], 
        lty = 2)
}

legend("topright", lwd = 2, lty = c(2, 1, 1, 1), col = c("black", "red", "blue", "green"), legend = c("Control", "4 cm", "8 cm", "16 cm"), bty = "n")


##### Model 1 residuals #####
plot(resid(glm.1) ~ fitted(glm.1), xlab="Fitted (Model 1)", ylab = "Residuals (Model 1)",
     main = "F. Residuals vs fitted (Model 1)", cex = 2)

##### Model 2 distribution #####

muphi.betaModel2 <- unique(data.frame(
  mu = fitted(glm.2),
  phi = predict(glm.2, type = "disp"),
  cardSize = shimData$cardSize,
  colourCards = shimData$colourCards,
  bgColour = shimData$bgColour
))

muphi.betaModel2.SUB <- muphi.betaModel2[muphi.betaModel2$colourCards == "B" & muphi.betaModel2$bgColour == "Gray",]


curve(dbeta2(x, muphi.betaModel2.SUB[1, 1], muphi.betaModel2.SUB[1, 2]),
      xlab = "proportion", ylab = "probability density",
      main = "B. Model 2. Hive ID as fixed effect, fixed phi"
)

abline(v = muphi.betaModel2.SUB[1, 1], lty = 2)

for (i in 1:4) {
  curve(dbeta2(x, muphi.betaModel2.SUB[i, 1], muphi.betaModel2.SUB[i, 2]), add = T, 
        col = c("blue", "black", "red", "green")[i], 
        lty = 1)
  abline(v = muphi.betaModel2.SUB[i, 1], col = c("blue", "black", "red", "green")[i], 
        lty = 2)
}

legend("topright", lwd = 2, lty = c(2, 1, 1, 1), col = c("black", "red", "blue", "green"), legend = c("Control", "4 cm", "8 cm", "16 cm"), bty = "n")


##### Model 2 residuals #####
plot(resid(glm.2) ~ fitted(glm.2), xlab="Fitted (Model 2)", ylab = "Residuals (Model 2)",
     main = "G. Residuals vs fitted (Model 2)", cex = 2)

##### Model 3 distribution #####

muphi.betaModel3 <- unique(data.frame(
  mu = fitted(glm.3),
  phi = predict(glm.3, type = "disp"),
  cardSize = shimData$cardSize,
  colourCards = shimData$colourCards,
  bgColour = shimData$bgColour
))

muphi.betaModel3.SUB <- muphi.betaModel3[muphi.betaModel3$colourCards == "B" & muphi.betaModel3$bgColour == "Gray",]


curve(dbeta2(x, muphi.betaModel3.SUB[1, 1], muphi.betaModel3.SUB[1, 2]),
      xlab = "proportion", ylab = "probability density",
      main = "C. Model 3. Hierarchical model, single phi"
)

abline(v = muphi.betaModel3.SUB[1, 1], lty = 2)

for (i in 1:4) {
  curve(dbeta2(x, muphi.betaModel3.SUB[i, 1], muphi.betaModel3.SUB[i, 2]), add = T, 
        col = c("blue", "black", "red", "green")[i], 
        lty = 1)
  abline(v = muphi.betaModel3.SUB[i, 1], col = c("blue", "black", "red", "green")[i], 
        lty = 2)
}

legend("topright", lwd = 2, lty = c(2, 1, 1, 1), col = c("black", "red", "blue", "green"), legend = c("Control", "4 cm", "8 cm", "16 cm"), bty = "n")


##### Model 3 residuals #####
plot(resid(glm.3) ~ fitted(glm.3), xlab="Fitted (Model 3)", ylab = "Residuals (Model 3)",
     main = "H. Residuals vs fitted (Model 3)", cex = 2)

##### Model 4 distribution #####

muphi.betaModel4 <- unique(data.frame(
  mu = fitted(glm.4),
  phi = predict(glm.4, type = "disp"),
  cardSize = shimData$cardSize,
  colourCards = shimData$colourCards,
  bgColour = shimData$bgColour
))

  muphi.betaModel4.SUB <- muphi.betaModel4[muphi.betaModel4$colourCards == "B" & muphi.betaModel4$bgColour == "Gray",]


curve(dbeta2(x, muphi.betaModel4.SUB[1, 1], muphi.betaModel4.SUB[1, 2]),
      xlab = "proportion", ylab = "probability density",
      main = "D. Model 4. Hierarchical model,\n variable phi (stimulus size)"
)

abline(v = muphi.betaModel4.SUB[1, 1], lty = 2)

for (i in 1:4) {
  curve(dbeta2(x, muphi.betaModel4.SUB[i, 1], muphi.betaModel4.SUB[i, 2]), add = T, 
        col = c("blue", "black", "red", "green")[i], 
        lty = 1)
  abline(v = muphi.betaModel4.SUB[i, 1], col = c("blue", "black", "red", "green")[i], 
        lty = 2)
}

legend("topright", lwd = 2, lty = c(2, 1, 1, 1), col = c("black", "red", "blue", "green"), legend = c("Control", "4 cm", "8 cm", "16 cm"), bty = "n")


##### Model 4 residuals #####
plot(resid(glm.4) ~ fitted(glm.4), xlab="Fitted (Model 4)", ylab = "Residuals (Model 4)",
     main = "I. Residuals vs fitted (Model 4)", cex = 2)

##### Model 5 distribution #####

muphi.betaModel5 <- unique(data.frame(
  mu = fitted(glm.5),
  phi = predict(glm.5, type = "disp"),
  cardSize = shimData$cardSize,
  colourCards = shimData$colourCards,
  bgColour = shimData$bgColour
))

  muphi.betaModel5.SUB <- muphi.betaModel5[muphi.betaModel5$colourCards == "B" & muphi.betaModel5$bgColour == "Gray",]


curve(dbeta2(x, muphi.betaModel5.SUB[1, 1], muphi.betaModel5.SUB[1, 2]),
      xlab = "proportion", ylab = "probability density",
      main = "E. Model 5. Hierarchical model,\n variable phi (stimulus shade)"
)

abline(v = muphi.betaModel4.SUB[1, 1], lty = 2)

for (i in 1:4) {
  curve(dbeta2(x, muphi.betaModel4.SUB[i, 1], muphi.betaModel4.SUB[i, 2]), add = T, 
        col = c("blue", "black", "red", "green")[i], 
        lty = 1)
  abline(v = muphi.betaModel4.SUB[i, 1], col = c("blue", "black", "red", "green")[i], 
        lty = 2)
}

legend("topright", lwd = 2, lty = c(2, 1, 1, 1), col = c("black", "red", "blue", "green"), legend = c("Control", "4 cm", "8 cm", "16 cm"), bty = "n")


##### Model 4 residuals #####
plot(resid(glm.4) ~ fitted(glm.4), xlab="Fitted (Model 4)", ylab = "Residuals (Model 4)",
     main = "J. Residuals vs fitted (Model 5)", cex = 2)

```

**Figure S3. A-E) Probability densities of shimmering response to black stimuli against a grey background predicted from the MLE parameters for Models 1-5.** *‘Control’* denotes the experimental condition where the string/wire alone was moved, and 4 cm, 8 cm and 16 cm are the diameters of the circular black stimuli. The *coloured lines* are density distributions of the shimmering response back-transformed for the sake of comparison with the original data. **(F-J)  Standardised residuals against the fitted values** The spread of residuals is plotted along the y-axis against the experimental treatments on the x-axis.

<br>
<br>
<br>

```{r betaModelTable1, echo=FALSE, message=FALSE, warning=FALSE}
### Beta-regression model AIC comparison
modelCompTable <- bbmle::AICtab(glm.0, glm.01, glm.1, glm.2, glm.3, glm.4, glm.5)
modelDetails <- c("Hierarchical, variable precision (stimulus size)", "Hierarchical, variable precision (stimulus shade)", "Hive identity as fixed effect", "Pooled-data, single precision", "Hierarchical, single precision", "Pooled-data null model", "Hierarchical null model")
modelCompTable$modelDetails <- modelDetails
names(modelCompTable) <- c("dAIC", "Degrees of freedom", "Model description")

knitr::kable(modelCompTable)
```

**Table S1. Comparison of the beta-regression models quantifying the shimmering response of black stimuli against grey background during bright light conditions**

```{r betaModelTable2, echo=FALSE, message=FALSE, warning=FALSE}
### Beta-regression model AIC comparison
ambCompTable <- bbmle::AICtab(glm.02, glm.03, glm.6, glm.7)
ambModelDetails <- c("Pooled-data model", "Hierarchical model", "Pooled-data null model", "Hierarchical null model")
ambCompTable$ambModelDetails <- ambModelDetails
names(ambCompTable) <- c("dAIC", "Degrees of freedom", "Model description")

knitr::kable(ambCompTable)
```
**Table S2. Beta-regression models comparing the shimmering response during daylight to that occurring in twilight**


\newpage

*This document was generated from the [supplementary markdown file](./sajesh_etal_2022_rev_ESM.Rmd). The details of the session and the packages used to generate this file are printed below*


***Session and packages info***
```{r sessionInfo, include=T, echo=F, results='markup'}
sessionDF = devtools::session_info()
sessionDF$platform

packageDF = devtools::package_info()
loadedPackages <- packageDF[packageDF$attached == T,] %>%
  select(package, ondiskversion, source)
print.data.frame(loadedPackages)
```

<br>
<br>
<br>

*The last section of the accompanying markdown file *[(sajesh_etal_2022_ESM.Rmd)](./sajesh_etal_2022_ESM.Rmd)  *contains the code for Figs. 2-4 in the main text. The plots can be generated by changing the value of the* `include` *argument in the corresponding code-chunk to* `TRUE`
```{r mainFigCode, include=FALSE, echo=FALSE}

###### fig2: faceted by bgColour and colourCards #####
### Defining the labels of the primary and secondary axes
fig2 <- ggplot(shimData, aes(cardSize, avgArea)) + 
  geom_rect(data = shimData[shimData$bgColour == "Black",],
            xmin = -Inf,xmax = Inf,
            ymin = -0.02,ymax = Inf, fill = "black") +
  geom_rect(data = shimData[shimData$bgColour == "Gray",],
            xmin = -Inf,xmax = Inf,
            ymin = -0.02,ymax = Inf, fill = "grey60") +
  geom_boxplot_pattern(aes(pattern = hiveID),
                       fill = "white",
                       pattern_spacing = 0.025,
                       pattern_density = 0.85,
                       pattern_fill = "grey70", 
                       color = "white") +
  scale_size_identity() +
  scale_x_discrete(labels = c("0" = "0[0]",
                              "4" = "4[1.6]",
                              "8" = "8[3.3]",
                              "16" = "16[6.5]")) +
  scale_alpha_manual(values=c(1,0.1)) +
    labs(x = "Stimulus diameter/\nvisual subtense (cm [deg])", y=TeX('$\\textit{S}_{trial}$'), 
         pattern = "Hive ID")+
    facet_grid(cols = vars(bgColour), 
               rows = vars(colourCards),
               labeller = labeller(hiveID = c("A" = "Hive A",
                                              "B" = "Hive B"),
                                   bgColour = c("Black"="Black background",
                                                "Gray"="Grey background"),
                                   colourCards = c("G1"="Light Grey",
                                                   "G3"="Dark Grey",
                                                   "B" = "Black"))) +
  theme(
    text = element_text(size=20),
    panel.background = element_rect(fill = "transparent"), # bg of the panel
    plot.background = element_rect(fill = "transparent", color = NA), # bg of the plot
    panel.grid.major = element_blank(), # get rid of major grid
    panel.grid.minor = element_blank(), # get rid of minor grid
    # legend.background = element_rect(fill = "grey90"), # get rid of legend bg
    # legend.box.background = element_rect(colour = NA, fill = "transparent"), # get rid of legend panel bg
    axis.title.y=element_text(angle=0, vjust = 0.5),
    axis.ticks.length.x = unit(0.1, "cm"),
    # axis.text.x = element_blank(),
    strip.text.y = element_blank(),
    strip.text.x = element_blank(),
    legend.position = "none"
    ) + 
  geom_point(data = subset(shimData, colourCards == "B"),
             aes(x = 0.75, y = 0.35), colour = "black", size = 10)+
  geom_point(data = subset(shimData, colourCards == "B"),
             aes(x = 0.75, y = 0.35), colour = "white", shape = 1, size = 10)+
  geom_point(data = subset(shimData, colourCards == "G1"),
             aes(x = 0.75, y = 0.35), colour = "gray90", size = 10)+
  geom_point(data = subset(shimData, colourCards == "G1"),
             aes(x = 0.75, y = 0.35), colour = "white", shape = 1, size = 10)+
  geom_point(data = subset(shimData, colourCards == "G3"),
             aes(x = 0.75, y = 0.35), colour = "gray60", size = 10)+
  geom_point(data = subset(shimData, colourCards == "G3"),
             aes(x = 0.75, y = 0.35), colour = "white", shape = 1, size = 10)
  

ggsave("fig2.eps", device = "eps", height = 7, width = 7)


##### fig3: twilight vs daylight boxplot #####
fig3 <- ggplot(ambLightData, aes(x = lightCond, y = avgArea, shape = lightCond)) +
  geom_rect(xmin = -Inf,xmax = Inf,
            ymin = -0.02,ymax = Inf, color = "grey60", fill = "grey60") +
  scale_shape_manual(values = c("Daylight" = 21,
                                "Twilight" = 23))+
  geom_boxplot_pattern(aes(pattern = hiveID),
                       fill = "white",
                       pattern_spacing = 0.01,
                       pattern_density = 0.85,
                       pattern_fill = "grey70", 
                       color = "black") +
  labs(x="Ambient light period", y = TeX('$\\textit{S}_{trial}$'))+
  # labs(x="Ambient light period", y = "Strength of\n shimmering", pattern = "Hive ID")+
  annotate("text", x=0.8, y=0.48, label="n=26", color = "black", size = 5)+
  annotate("text", x=1.2, y=0.29, label="n=10", color = "black", size = 5)+
  annotate("text", x=1.8, y=0.17, label="n=25", color = "black", size = 5)+
  annotate("text", x=2.2, y=0.26, label="n=20", color = "black", size = 5)+
  theme(
    text = element_text(size=20),
    panel.background = element_rect(fill = "transparent"), # bg of the panel
    plot.background = element_rect(fill = "transparent", color = NA), # bg of the plot
    panel.grid.major = element_blank(), # get rid of major grid
    panel.grid.minor = element_blank(), # get rid of minor grid
    # legend.background = element_rect(fill = "grey90"), # get rid of legend bg
    # legend.box.background = element_rect(colour = NA, fill = "transparent"), # get rid of legend panel bg
    axis.title.y=element_text(angle=0, vjust = 0.5),
    axis.ticks.length.x = unit(0.1, "cm"),
    # axis.text.x = element_blank(),
    strip.text.y = element_blank(),
    strip.text.x = element_blank(),
    legend.position = "none"
    )+
  guides(shape = "none")

ggsave("fig3.eps", device = "eps", height = 7, width = 7)

##### Habituation plot #####
fig4a <- shimData %>%
  filter(colourCards == "B",
         cardSize == "16" | cardSize == "8",
         bgColour == "Gray") %>%
  mutate(timediff = ((trialTime - lag(trialTime))),
         carddiff = as.numeric(as.character(cardSize)) - lag(as.numeric(as.character(cardSize)))) %>%
  select(avgArea, hiveID, cardSize, trialTime, timediff, carddiff) %>%
  filter(timediff < 720) %>%
  ggscatter(x = "timediff", y = "avgArea", add = "reg.line", shape = "cardSize") + 
  stat_cor(label.y = 0.55, size = 5) + stat_regline_equation(label.y = 0.5, size = 5) + 
  labs(x = "Time elapsed since\n previous trial (min)", y=TeX('$\\textit{S}_{trial}$')) +
  facet_wrap(~hiveID, labeller = labeller(hiveID = c("A" = "Hive A",
                                                     "B" = "Hive B"))) + 
  theme(
    text = element_text(size=20),
    panel.background = element_rect(fill = "transparent"), # bg of the panel
    plot.background = element_rect(fill = "transparent", color = NA), # bg of the plot
    panel.grid.major = element_blank(), # get rid of major grid
    panel.grid.minor = element_blank(), # get rid of minor grid
    legend.background = element_rect(fill = "grey90"), # get rid of legend bg
    legend.box.background = element_rect(colour = NA,
                                         fill = "transparent"), # get rid of legend panel bg
    panel.spacing.x = unit(2, "lines"),
    axis.title.y=element_text(angle=0, vjust = 0.5),
    axis.ticks.length.x = unit(0.1, "cm"),
    # axis.text.x = element_blank(),
    strip.text.y = element_blank(),
    # strip.text.x = element_blank(),
    # legend.position = "none"
  )+
  guides(shape = "none")

fig4b <- shimData %>%
  filter(colourCards == "B",
         cardSize == "8" | cardSize == "16",
         bgColour == "Gray") %>%
  ggscatter(x = "trialTimes", y = "avgArea", add = "reg.line", shape = "cardSize") + 
  stat_cor(label.y = 0.55, size = 5) + stat_regline_equation(label.y = 0.5, size = 5) + 
  labs(x= "Time of trial (hh:mm)", y = TeX('$\\textit{S}_{trial}$')) + 
  facet_grid(~hiveID,
             labeller = labeller(.cols = c("A" = "Hive A",
                                           "B" = "Hive B"),
                                 .rows = c("8" = "8 cm",
                                           "16" = "16 cm"))) +
  theme(
    text = element_text(size=20),
    panel.background = element_rect(fill = "transparent"), # bg of the panel
    plot.background = element_rect(fill = "transparent", color = NA), # bg of the plot
    panel.grid.major = element_blank(), # get rid of major grid
    panel.grid.minor = element_blank(), # get rid of minor grid
    legend.background = element_rect(fill = "grey90"), # get rid of legend bg
    legend.box.background = element_rect(colour = NA,
                                         fill = "transparent"), # get rid of legend panel bg
    axis.title.y=element_text(angle=0, vjust = 0.5),
    axis.ticks.length.x = unit(0.1, "cm"),
    panel.spacing.x = unit(2, "lines"),
    strip.text.y = element_blank(),
    plot.margin = unit(c(0,0.75,0,0), units = "cm")
  )+
  guides(shape = "none")

fig4 <- cowplot::plot_grid(fig4a, fig4b, labels = "AUTO", ncol = 1, label_size = 20)

ggsave("fig4.eps", device = "eps", height = 9, width = 10)

```
````
